# Chronic Disease Emergence in Coevolving Social Networks via Behavioral Feedback and Demographic Turnover

**DOI:** 10.64898/2026.07.09.737414

**Authors:** Hsuan-Wei Lee, Ji-Yuan Lo, Martin O’Flaherty

**Affiliations:** Department of Biostatistics and Health Data Science, Lehigh University, Bethlehem, PA 18015, USA; Department of Finance, National Taiwan University, Taipei, Taiwan; Department of Public Health, Policy and Systems, Institute of Population Health, University of Liverpool, Liverpool, UK

**Keywords:** social networks, cardiovascular disease, agent-based model, coevolving networks, behavioral contagion, demographic turnover, network epidemiology, approximate Bayesian computation

## Abstract

Chronic diseases emerge within social systems where individuals reorganize their ties, adapt their behaviors, and turn over through demographic processes; yet, how these coupled dynamics shape long-run disease trajectories remains poorly understood. We developed an agent-based model of how the network structure, behavioral adaptation, and population renewal jointly generate cardiovascular disease incidence in an evolving population. The model integrated age-dependent intrinsic risk from the Framingham Risk Score framework, behavioral transmission through social ties, homophily-driven network formation, and demographic turnover across simulated populations of up to 420 agents and horizons of 18–240 months. Highly connected agents showed lower simulated incidence than weakly connected agents (adjusted hazard ratio 0.43; 95% confidence interval 0.33, 0.57; top-tertile degree indicator) after adjustment for age, sex, smoking, blood pressure, body-mass index, physical activity, and diet. Individual disease probability rose with ego-network disease prevalence (Pearson’s *r* = 0.703, *p* < 0.001). Global sensitivity analysis identified baseline incidence, activity level, and behavioral transmission as dominant drivers (78% of variance). Bayesian calibration concentrated within the empirical acceptance region, with posterior standard deviation of 12% of prior. These results point toward a systems-level understanding of non-communicable disease emergence.

## 1 Introduction

Chronic diseases do not develop in isolation. They emerge within social systems where individuals continuously interact, form and dissolve ties, adapt their behaviors in response to what they observe around them, and turn over through demographic processes. These coupled dynamics operate across timescales ranging from months to decades, and their joint effects on long-run disease trajectories remain poorly understood mechanistically. Cardiovascular disease (CVD) offers a compelling context to investigate these dynamics. It is the leading cause of mortality in the United States, affecting about 130.6 million adults (48.9%) and accounting for 915,973 deaths in 2023 [1]. Although CVD mortality has declined substantially over recent decades [2], these improvements have decelerated in several high-income countries and now contribute notably to the slowdown in life-expectancy gains [3]; trends also remain heterogeneous across demographic groups [4], suggesting that factors beyond conventional clinical risk alone shape the population-level patterns.

Social relationships constitute a well-established determinant of cardiovascular health. Meta-analytic evidence shows that social isolation increases the risk for coronary heart disease and stroke [5], while poor social relationships increase the overall mortality risk by approximately 50% [6]. These effects are comparable in magnitude to established risk factors such as smoking and obesity [7]. Prospective cohort studies have documented that socially isolated individuals face nearly a twofold increase in CVD mortality [8], and longitudinal analyses show that declining network size predicts an elevated cardiovascular risk independent of the baseline health status [9, 10]. Network position shapes health outcomes through multiple pathways [11, 12], including stress buffering, access to health information [13], and the spread of health behaviors through social ties, with large cohort studies documenting the clustering of obesity and smoking within social networks [14, 15]. Yet, associational evidence alone cannot reveal the mechanisms by which network structure and disease risk coevolve over time.

Empirical studies examining the network effects on health face fundamental methodological constraints. Homophily, the tendency to associate with similar others [16], can produce network clustering of disease that superficially resembles contagion but reflects selection rather than influence [17]. Disentangling these processes requires longitudinal network data with repeated measures of both ties and behaviors, which remain scarce for chronic diseases with long latency periods [18–20]. Observational studies also struggle to establish the temporal ordering between network change and health outcomes, since individuals may alter their social ties in response to illness [21]. Network characteristics such as density and centrality have been associated with health outcomes in diverse populations [22–24], although causal interpretation remains difficult without experimental or quasi-experimental designs. These constraints motivate a complementary computational approach that can make selection and influence explicit, separate them mechanistically, and study their joint effects over long timescales.

Agent-based models provide exactly this capability. By simulating populations of heterogeneous individuals whose behaviors and network ties evolve according to explicit rules, they allow systematic investigation of scenarios that observational studies cannot access [25, 26]. Previous work has shown that disease transmission and network topology coevolve dynamically, with social clustering amplifying the spread, while networks reorganize to reduce exposure [27, 28]. Behavioral feedback in epidemic models suggests that awareness of disease risk can alter contact patterns and dampen transmission [29, 30]. Population-level models have successfully decomposed historical cardiovascular mortality trends and projected future burden under alternative policy scenarios [2, 31]. However, many population-level models treat individuals as independent units and therefore cannot represent how behavioral change propagates through evolving social ties [32, 33]. Extending coevolving network frameworks to chronic disease requires a different dynamical structure. CVD is represented here as an absorbing state, so affected agents do not return to a susceptible state. Risk accumulates gradually over long time horizons rather than resolving during the short timescales typical of acute infection. Demographic turnover also renews the population while altering the network context in which future disease risk unfolds [34, 35]. Together, these features create feedback structures that standard epidemic models were not designed to represent.

We developed an agent-based model that couples network coevolution, behavioral adaptation, and demographic turnover to examine how chronic disease trajectories emerge in adaptive social systems. The model assigns each agent an intrinsic CVD risk derived from the Framingham Risk Score framework [36, 37], which is modulated by behavioral transmission through social ties and the homophily-driven network formation. Population composition evolves continuously through agent entry and exit. We used the model to address three questions: whether persistent differences in disease trajectories emerge across network positions after adjustment for individual-level risk factors, whether local exposure generates dose-response patterns in simulated disease probability, and how demographic turnover and behavioral adaptation jointly shape long-run prevalence at the population level.

## 2 Methods

### 2.1 Model overview

We developed an agent-based model to study how CVD incidence evolves in a dynamic social network with behavioral adaptation and population turnover. The simulated population was of a fixed size with *N* agents. At each monthly time step, agents could age, develop CVD, adjust their social activity, form or dissolve ties, and exit the population. Exiting agents were immediately replaced by new entrants, keeping the total population size constant while allowing the age composition and disease prevalence to evolve. The model was implemented in Python using `NetworkX` for graph operations, with fixed random seeds to ensure reproducibility, and it integrated three coupled components: an individual disease-risk module, a dynamic social-network module, and a population-turnover module. This framework is a hypothesis-generating theoretical tool rather than a predictive model of real-world CVD incidence; parameter values were chosen to produce dynamically plausible behavior rather than estimated from longitudinal data.

### 2.2 Agent attributes

Each agent *i* was initialized with demographic, behavioral, and physiological attributes. Age was sampled from three categories: 18–35, 36–55, and 55+ years, with a continuous uniform draw within each category. Sex was assigned as male with probability 0.49 and female with probability 0.51. A latent health-orientation score, *h*_*i*_, governed correlated behavioral and clinical attributes and was drawn as

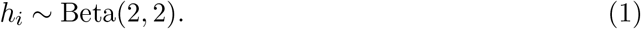

Agents with higher *h*_*i*_ tended toward healthier behavioral profiles. Attributes governed by this score included smoking status (never, former, or current), physical activity (low, moderate, or high), diet quality (poor, average, or healthy), body-weight category (underweight, normal, overweight, or obese), and blood-pressure category (normal, pre-hypertension, or hypertension). Agents also differed in three behavioral-response parameters governing how strongly they reacted to perceived disease risk. Fear sensitivity was drawn from *N* (1.0, 0.2) and clipped to [0.5, 1.6]. Response elasticity was drawn from *N* (1.0, 0.25) and clipped to [0.5, 1.8]. The ego-network weighting parameter, *ω*_*i*_, determined how much weight each agent placed on their local neighborhood relative to the population average when forming disease-risk perceptions; it was drawn from *N* (0.35, 0.08) and clipped to [0.15, 0.55], unless a specific analysis fixed this parameter to isolate the ego-network perception.

### 2.3 CVD onset probability

Each month, healthy agents faced a combined CVD onset probability integrating intrinsic risk and social transmission:

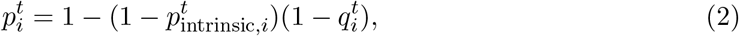

clipped to [0, 0.95]. The two components are described in turn.

The intrinsic risk component captured individual biological and behavioral vulnerability, following the multiplicative structure of the Framingham Risk Score framework [36, 37]:

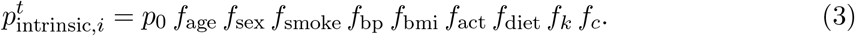

Here, *p*_0_ is the baseline monthly incidence rate. The age multiplier increased linearly above age 40,

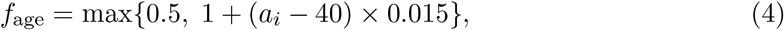

and was bounded below at 0.5 to prevent implausibly low values in young agents. The sex multiplier was 1.4 for male agents and 1.0 for female agents, consistent with epidemiological evidence on sex differences in CVD incidence [38]. Risk multipliers for behavioral and clinical attributes were calibrated to published relative-risk estimates: 1.0, 1.6, and 2.7 for never, former, and current smokers, respectively; 1.0, 1.5, and 2.0 for normal blood pressure, pre-hypertension, and hypertension, respectively; 1.0, 1.0, 1.4, and 1.8 for underweight, normal, overweight, and obese body weight, respectively; 1.0, 0.85, and 0.75 for low, moderate, and high physical activity, respectively; and 1.0, 0.8, and 0.70 for poor, average, and healthy diet, respectively.

These behavioral and clinical multipliers were anchored to large pooled or trial estimates rather than chosen freely. The current-smoker multiplier (2.7) reflects the elevated acute myocardial infarction risk reported in the INTERHEART study, in which current smoking carried an odds ratio of approximately 2.9 [39]; a slightly more conservative value was adopted here because that estimate is specific to acute myocardial infarction and would overstate the risk for the broader composite cardiovascular outcome represented in this model. The high physical-activity multiplier (0.75) corresponds to the roughly 20–30% reduction in cardiovascular risk associated with higher leisure-time activity in the pooled cohort evidence [40]. The healthy-diet multiplier (0.70) reflects the approximately 30% relative reduction in major cardiovascular events observed for a Mediterranean dietary pattern in the PREDIMED trial [41]. Intermediate categories were set to interpolate between these anchors. Because the model is multiplicative and operates on a monthly baseline hazard, these values were intended to reproduce realistic relative gradients rather than recover absolute trial effect sizes.

Network position additionally modified intrinsic risk through baseline degree *k*_0_ and local clustering coefficient *c*_0_, both measured after a network burn-in period:

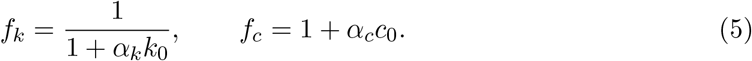

Higher degree reduced the intrinsic risk through *f*_*k*_, reflecting the buffering effect of social connectivity. Higher local clustering slightly elevated risk through *f*_*c*_, reflecting greater exposure within tightly connected neighborhoods. CVD status was absorbing: once an agent developed CVD, it remained CVD positive for the remainder of the simulation. This absorbing-state assumption distinguishes the model from infectious disease frameworks where recovery is possible and may produce different long-run patterns of network-disease coevolution.

The social-transmission component captured behavioral contagion through network exposure, combining two channels: the count of CVD-positive neighbors and the weighted frequency of contact with them:

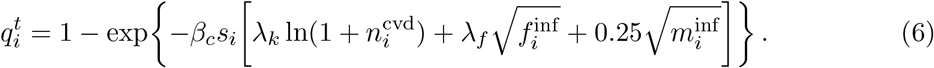

Here, *β*_*c*_ is the contagion-strength parameter, 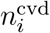 is the number of CVD-positive neighbors, 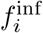 is the infected contact frequency, and 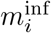 is a memory-decayed signal of past infected contact. The logarithmic form for neighbor count reflects diminishing marginal returns to exposure breadth. The square-root form for contact frequency reflects sub-linear scaling with exposure intensity. Individual susceptibility was

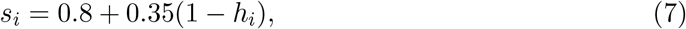

with an additional 0.08 added for current smokers. Unless otherwise specified, (*λ*_*k*_, *λ*_*f*_) = (0.55, 0.45). Each agent aged by 1*/*12 year at every monthly step.

### 2.4 Behavioral feedback and social exposure

Agents formed a perceived CVD prevalence by blending population-level information with ego-network experience, updated at each step as

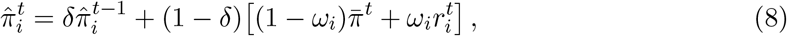

where 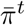 is population prevalence, 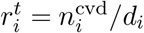 is the proportion of CVD-positive neighbors, *d*_*i*_ is degree, and *δ* = 0.72 is a memory-decay parameter that smoothed perceived prevalence over time. Agents with higher *ω*_*i*_ weighted their local neighborhood more heavily than the global signal.

Perceived prevalence then fed back into social activity. Agents reduced their activity as perceived risk rose, according to

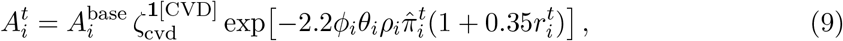

clipped to [*η*_floor_, 1.0] with *η*_floor_ = 0.02. Here, *ϕ*_*i*_ is behavioral feedback strength, *θ*_*i*_ is fear sensitivity, *ρ*_*i*_ is response elasticity, and *ζ*_cvd_ is a multiplicative reduction applied to agents who had already developed CVD. Tie strength between agents *u* and *v* depended on their joint activity level and pairwise similarity:

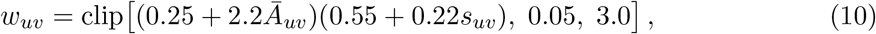

where Ā_*uv*_ is the mean activity of the dyad. Infected contact frequency summed weighted exposure across CVD-positive neighbors:

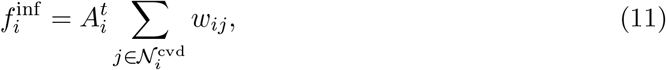

where 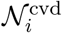 denotes the CVD-positive neighbors of agent *i*.

### 2.5 Dynamic social network

The initial network was constructed as a Barabási-Albert preferential-attachment graph with *m* = 3 edges per new node, producing a scale-free degree distribution with a hub structure. This was followed by 4*N* triadic-closure enhancement rounds in which two random neighbors of a pivot node were connected with probability 0.7, increasing local clustering toward empirically realistic levels [42]. Pairwise similarity between agents *u* and *v* combined the demographic, behavioral, and disease-status components:

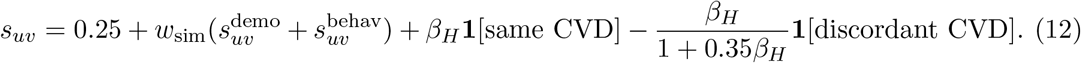

Demographic similarity 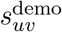 included age group, sex, smoking status, and diet quality, with weights 0.80, 0.35, 0.35, and 0.25, respectively. Parameter *β*_*H*_ governed CVD-based homophily, with higher values causing CVD-concordant pairs to form ties preferentially.

At each monthly step, the network evolved through three mechanisms. Existing edges survived independently with probability *π*_edge_, and surviving ties decayed in strength at rate *δ*_*e*_. New edges were then proposed through *M* candidate-pair attempts, of which a fraction, *p*_triad_, was selected via triadic closure. For each candidate pair (*u, v*) without an existing tie, formation occurred with probability

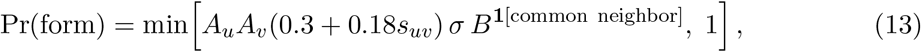

where *σ* is a formation-scale parameter and *B* is a triadic-closure boost applied when the pair shared at least one common neighbor. Default network parameters were *π*_edge_ = 0.95, *δ*_*e*_ = 0.02, *σ* = 0.40, *p*_triad_ = 0.40, and *B* = 3.0.

### 2.6 Population turnover

Agents exited the population each month according to an age-dependent and disease-dependent hazard. The monthly exit probability was

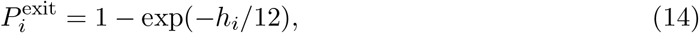

where the annualized exit hazard was

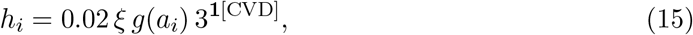

with exit multiplier *ξ* and age-scaling function

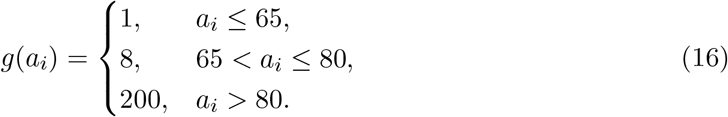

CVD-positive agents faced a threefold elevated exit rate, reflecting elevated mortality. Older agents faced sharply higher exit rates through *g*(*a*_*i*_). Each exiting agent was immediately replaced by a new entrant drawn from the specified entry age distribution, preserving population size while allowing age composition and disease prevalence to evolve dynamically. Population renewal continuously introduced younger healthy agents, acting as a natural dampening mechanism on long-run prevalence.

### 2.7 Monthly update sequence

At each monthly step the model updated in a fixed sequence. Tie strengths were refreshed, population prevalence and ego-network exposure were computed, and agents updated their perceived prevalence and social activity. CVD onset was then evaluated for each healthy agent, and all agents aged by one month. Edges were retained or dissolved, surviving ties decayed, and new ties formed through homophily, activity-dependent mixing, and triadic closure, after which population turnover was applied. Disease risk at month *t* therefore depended on network exposure before turnover, while rewiring and replacement determined the structure available at month *t* + 1.

### 2.8 Simulation design

All analyses used the same model architecture and update rules. Parameter settings varied only to isolate specific mechanisms or evaluate robustness, and they are documented in full in Supplementary Table S1.

For population-dynamics analyses, the model used *N* = 300 agents simulated over 150–240 months. The initial age distribution placed 30% of agents in the 18–35 group, 40% in the 36–55 group, and 30% in the over-55 group. Replacement entrants were drawn from a younger distribution (75%, 23%, and 2% across the same groups) to produce realistic demographic renewal.

For survival analyses, simulations used *N* = 420 agents over 120 months. Degree-stratified analyses used a 24-month network burn-in, 2,200 edge-formation attempts per month, *α*_*k*_ = 0.15, and *α*_*c*_ = 0.60. Baseline degree was grouped into tertiles. Multivariable survival models pooled six independent replicates and adjusted for age, sex, obesity, network clustering, physical activity, diet quality, smoking status, blood pressure, and network degree.

For homophily and social-contagion analyses, the model used *N* = 300 agents with a 24-month burn-in. The base configuration used *p*_0_ = 0.0003, *β*_*c*_ = 0.08, *β*_*H*_ = 6.0, *π*_edge_ = 0.97, *σ* = 0.9, *p*_triad_ = 0.5, *B* = 4.0, *α*_*k*_ = 0.05, *α*_*c*_ = 2.0, exit rate 0.008, behavioral feedback 0.10, and *ζ*_cvd_ = 0.85. Homophily comparisons contrasted *β*_*H*_ = 6.0 with *β*_*H*_ = 0.0. Dose-response analyses recorded the proportion of CVD-positive neighbors and own CVD status across repeated network snapshots.

Behavioral-feedback analyses used *N* = 340 *− −*380 agents over 150–160 months with a 24-month burn-in. In the activity-feedback analysis, *ω*_*i*_ was fixed at 1.0 so that perceived prevalence reflected only ego-network exposure. Two-dimensional parameter sweeps used *N* = 220, *T* = 110 months, an 18-month burn-in, *p*_0_ = 0.0016, *β*_*H*_ = 6.2, exit rate 0.008, and five replicates per grid cell. One sweep varied *π*_edge_ *∈* [0.75, 0.99] and *σ ∈* [0.20, 0.80] on an 11 × 11 grid. A second sweep varied behavioral feedback strength *ϕ ∈* [0.0, 1.2] and CVD exit multiplier *ξ ∈* [1.0, 6.5] on an 11 × 11 grid. Raw cell means were blended with a linear trend surface fit by ordinary least squares for visualization.

### 2.9 Global sensitivity analysis

We used Latin hypercube sampling to assess how output variance was distributed across eight model parameters: baseline incidence rate *p*_0_ *∈* [10^*−*5^, 3 × 10^*−*4^], homophily boost *β*_*H*_ *∈* [0.5, 5.0], behavioral transmission rates in [0.10, 1.50] (scaling *λ*_*k*_ and *λ*_*f*_ proportionally), exit multiplier *ξ ∈* [1.0, 4.0], edge decay *δ*_*e*_ *∈* [0.0, 0.08], memory decay *δ ∈* [0.30, 0.90], similarity weight *w*_sim_ *∈* [0.20, 1.50], and activity scale in [0.60, 1.20]. Thirty-six parameter sets were drawn by Latin hypercube sampling. Each was evaluated with four independent replicates using *N* = 50 agents, *T* = 18 months, and a six-month burn-in. Partial rank correlation coefficients were computed from rank-transformed input and output matrices, with bootstrap standard errors from 300 resamples. Cumulative parameter importance was computed from squared partial rank correlation coefficients normalized to sum to 100%.

### 2.10 Bayesian calibration

We used approximate Bayesian computation with rejection sampling as a calibration check, targeting an empirical CVD prevalence of 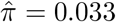 derived from published epidemiological estimates. Prior samples were drawn by Latin hypercube sampling over four parameters: incidence rate, behavioral transmission rate, edge decay, and exit multiplier, using the same ranges as the sensitivity analysis. A total of 120 prior samples were evaluated with three independent replicates each, using *N* = 50 agents and *T* = 18 months. Samples were accepted when mean final prevalence fell within the interval [0.000, 0.073] around the target. Posterior medians and 95% credible intervals were computed from accepted samples. Prior and posterior predictive distributions were compared to assess whether calibration concentrated simulated prevalence near the empirical target.

### 2.11 Statistical analysis

Kaplan-Meier curves estimated CVD-free survival across age and network-degree groups. Cox proportional-hazards models were estimated with a ridge penalty of *λ* = 0.01 when multivariable adjustment was required. Group differences in survival were assessed with log-rank tests. The primary exposure in degree-stratified analyses was the baseline network degree grouped into tertiles. Dose-response relationships were evaluated using Pearson correlation, Spearman rank correlation, and ordinary least squares regression, as appropriate. Locally estimated scatterplot smoothing was used for descriptive curves, with bootstrap confidence bands for smoothed degree-risk relationships. Partial rank correlation coefficients were used for global sensitivity analysis. All analyses were conducted in Python 3 using `NumPy, pandas, NetworkX, lifelines, statsmodels, scipy,` and `matplotlib.`

### 2.12 Model validation

Because datasets combining repeated social-network measurements with chronic disease incidence over extended follow-up remain scarce, direct validation against longitudinal network data was not possible. Validation therefore focused on internal consistency, face validity, and calibration to empirically supported prevalence targets. We verified that intrinsic risk increased with age, that rising perceived prevalence suppressed social activity, and that network statistics and limiting-case behavior remained plausible, confirming that the model behaved consistently with its stated assumptions before substantive analyses were conducted.

## 3 Results

We first characterized system-level dynamics emerging from demographic turnover and network adaptation. We then examined whether persistent differences arose across network positions, investigated mechanisms underlying these patterns, and evaluated model robustness and parameter identifiability.

### 3.1 Population dynamics and network evolution

Population turnover generated a natural dampening effect on long-run disease burden. Without replacement, prevalence in the original cohort rose monotonically over 240 months, whereas replacing exiting agents with younger healthy entrants held total population preva-lence near its 120-month level. The two trajectories diverge progressively, showing that demographic turnover is itself a structural determinant of population-level disease burden, independent of any change in individual-level risk. Age distributions behaved correspondingly, with younger replacement cohorts producing intermediate age compositions relative to the closed population, and network degree declined gradually to approximately 3.4 after 150 months (Supplementary Figure S1).

**Figure 1:**
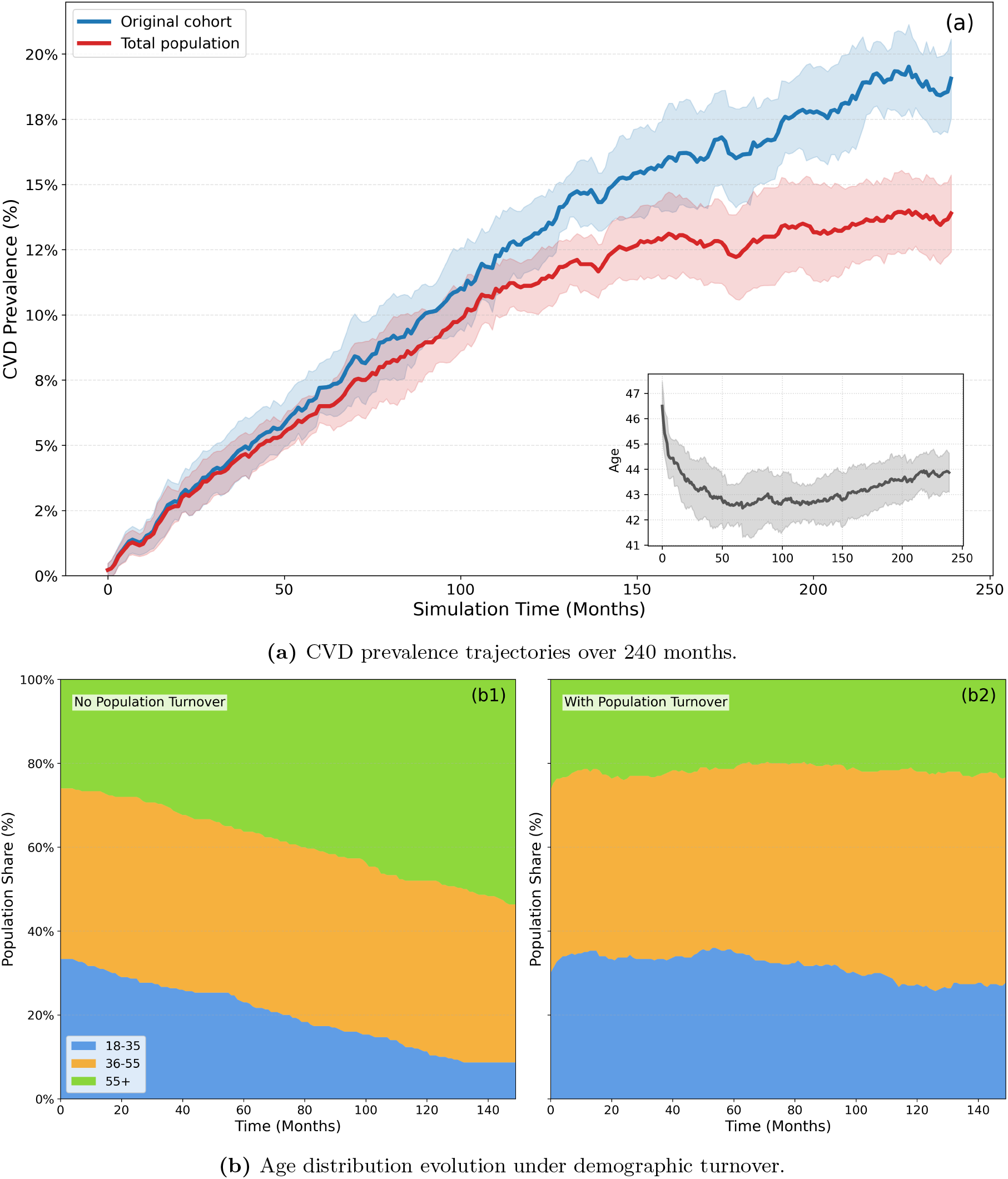
Population dynamics under demographic turnover. Panel A shows that cardiovascular disease (CVD) prevalence in the original cohort (blue line) rose to approximately 19% over 240 months, while total population prevalence (red line) stabilized near 14% after 120 months, reflecting the equilibrium between incidence in aging agents and replacement by healthy young entrants. Panel B shows the age distribution evolution: without turnover, the population aged monotonically, whereas younger replacement cohorts produced intermediate age distributions. Network degree evolution is presented in Supplementary Figure S1.

### 3.2 Survival analysis stratified by network position

Kaplan-Meier curves stratified by age reproduced known epidemiological gradients, providing a face-validity check for the intrinsic risk module, with agents initially aged 55 and above experiencing substantially faster CVD onset than younger groups.

Network degree produced a distinct and persistent effect on simulated survival that was not attributable to age or individual risk factors. Agents in the high-degree tertile (degree *≥* 5) showed better CVD-free survival throughout the observation period than agents in the low-degree tertile (degree *≤* 2), with the two curves diverging progressively over time. Cox proportional hazards regression adjusting for age, sex, smoking status, blood pressure category, body-mass index category, physical activity level, and diet quality yielded an adjusted hazard ratio (HR) of 0.43 (95% confidence interval [CI] 0.33–0.57) for the top-tertile degree indicator. In the degree-stratified survival model, the high-degree tertile had HR 0.59 (95% CI 0.43–0.81) relative to the low-degree tertile. Older agents showed elevated HRs (age 55+ versus 18–35: HR 2.08, 95% CI 1.60–2.70). Current smoking, hypertension, and obesity were each associated with elevated simulated hazard, consistent with their known epidemiological roles [38].

**Figure 2:**
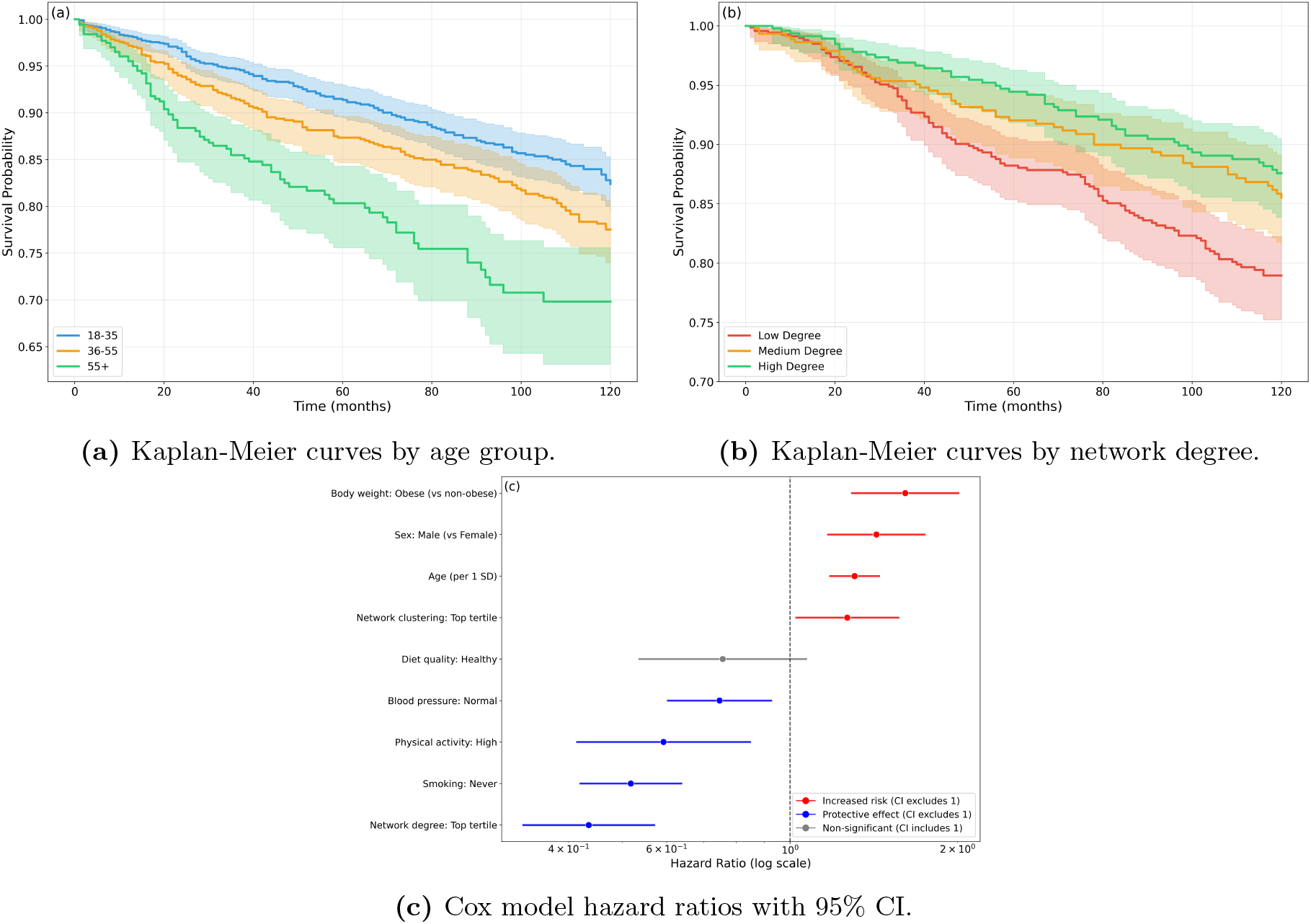
Simulated survival differences by age group and network degree. Panel A shows that age gradients in cardiovascular disease (CVD)-free survival were consistent with known epidemiological patterns, providing a face-validity check for the intrinsic risk module. Panel B shows that highly connected agents consistently exhibited better CVD-free survival than weakly connected agents, with survival curves diverging progressively over 120 months. Panel C presents Cox model hazard ratios adjusted for age, sex, smoking status, blood pressure, body-mass index, physical activity, and diet quality. The adjusted hazard ratio for the top-tertile degree indicator was 0.43 (95% confidence interval [CI]: 0.33–0.57). Forest plot point colors indicate the direction and significance of each association.

### 3.3 Homophily, social contagion, and network structure

Comparing strong homophily (*β*_*H*_ = 6.0) with no homophily (*β*_*H*_ = 0.0) showed that the degree-CVD relationship in this experiment was dominated by saturation: probability rose with degree and approached one under no homophily, whereas strong homophily produced an abrupt low-to-high transition over a narrow degree range. This panel therefore reflects saturation and clustering under the homophily-contagion regime rather than a stable protective or smooth monotone degree effect.

**Figure 3:**
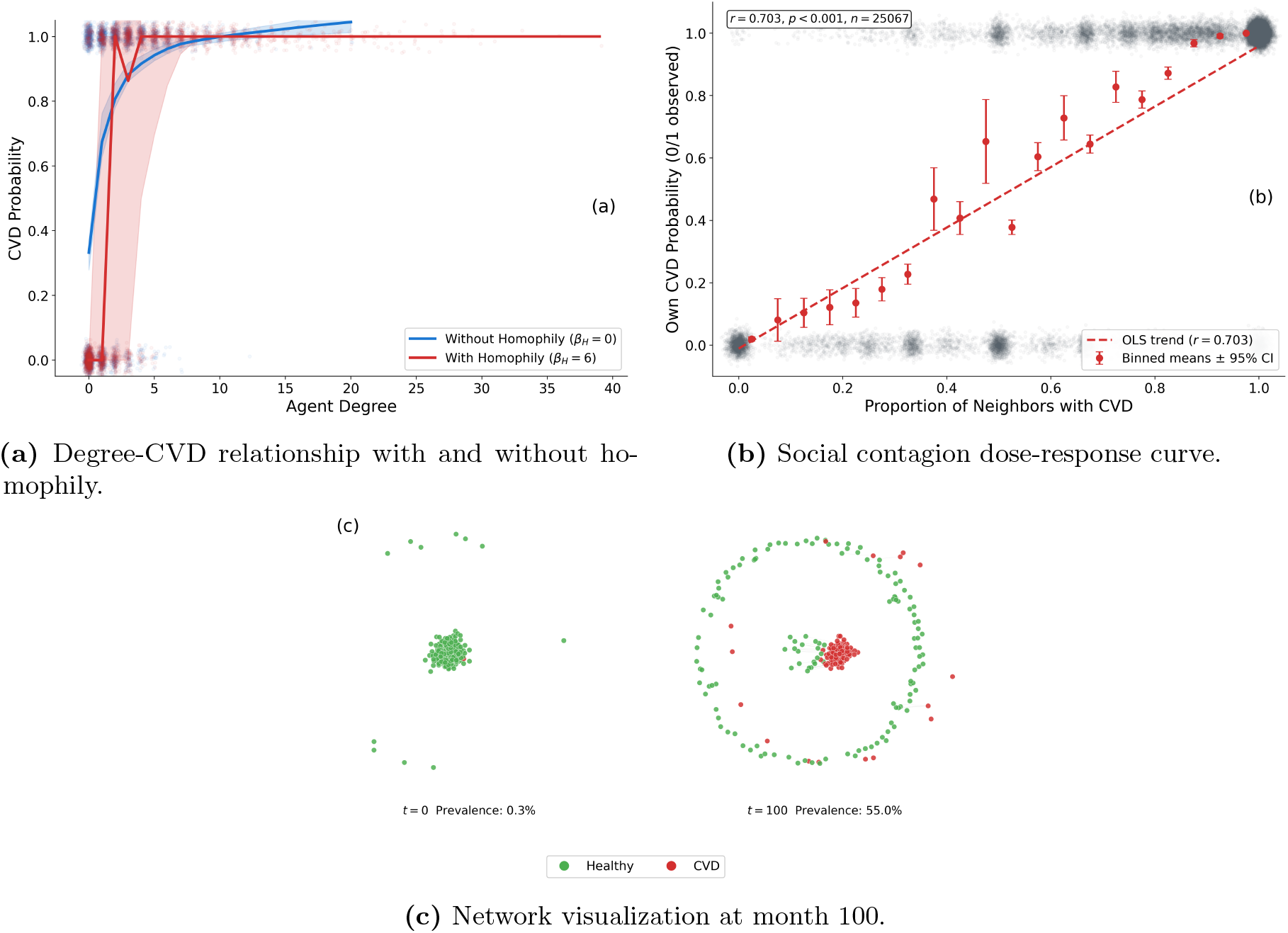
Homophily and dose-response patterns in simulated disease transmission. Panel A shows that, under the homophily-contagion regime used for this analysis, cardiovascular disease (CVD) probability rapidly approached saturation across much of the observed degree range. The no-homophily curve rose with degree before saturating, whereas the homophily condition showed an abrupt transition with wide uncertainty at low degrees, consistent with near-binary separation rather than a smooth protective degree gradient. Panel B shows a strong dose-response relationship between the proportion of CVD-positive neighbors and individual CVD probability (Pearson *r* = 0.703, *p <* 0.001), consistent with behavioral transmission through network ties. Panel C illustrates the network structure at month 100, with CVD-positive agents (red) showing moderate spatial clustering consistent with homophily-driven assortative mixing alongside ongoing random edge formation.

Individual CVD probability increased monotonically with the proportion of CVD-positive neighbors (Pearson *r* = 0.703, *p <* 0.001), a strong dose-response pattern. Because the model incorporated both homophilic selection and network influence as explicit mechanisms, this pattern reflected the joint operation of both processes and should not be interpreted as a purely causal transmission effect. Network visualizations showed scale-free initial structure with clear hubs, and by month 100 CVD-positive agents exhibited moderate spatial clustering rather than complete segregation, consistent with homophily-driven assortative mixing alongside ongoing random edge formation.

### 3.4 Global sensitivity analysis

Sensitivity analysis identified a clear hierarchy of parameters governing simulated CVD prevalence. Baseline incidence rate *p*_0_ was the dominant driver (PRCC = +0.80), with activity rates and behavioral transmission rates forming a closely ranked second tier (both PRCC = +0.52), followed by the exit multiplier (PRCC = *−*0.42), the only negatively signed parameter, and homophily boost (PRCC = +0.33). Similarity weight, edge decay, and memory decay contributed little (PRCC = +0.15, +0.14, and +0.03), with uncertainty ranges spanning zero. The top-ranked parameter alone explained about 42% of output variance and the top three explained 78.2%, with the cumulative curve reaching roughly 97% by rank five. This concentration indicates a parsimonious structure in which the dominant mechanisms are identifiable and interpretable.

**Figure 4:**
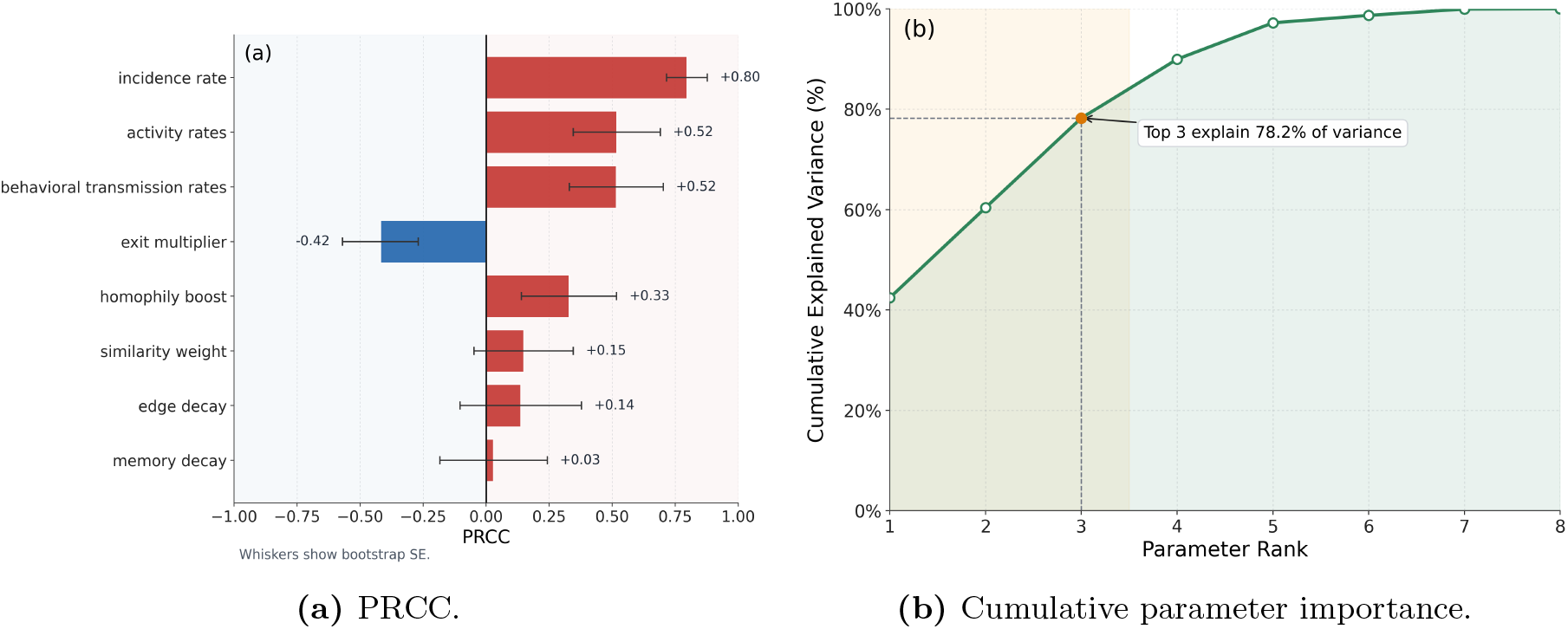
Global sensitivity analysis identifies dominant drivers of model output. Panel A shows the partial rank correlation coefficients (PRCCs) for eight parameters. Baseline incidence rate (*p*_0_) was the dominant driver (PRCC = +0.80). Activity rates and behavioral transmission rates formed a second tier with similar magnitudes (PRCC *≈* +0.52). The exit multiplier was the only strongly negatively signed parameter (PRCC = *−*0.42), reflecting the prevalence-dampening effect of population turnover. Panel B shows that the top three parameters jointly accounted for 78.2% of output variance, and the cumulative curve flattened substantially beyond rank five, indicating a parsimonious model structure.

### 3.5 Bayesian calibration

Approximate Bayesian computation accepted 71 of 120 prior samples (acceptance rate 59.2%). Calibration used a nominal prevalence target of 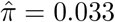, yielding an acceptance interval of [0.000, 0.073]. The posterior predictive mode was approximately 0.012, which fell within this acceptance interval; thus, the target and posterior peak refer to distinct quantities.

**Figure 5:**
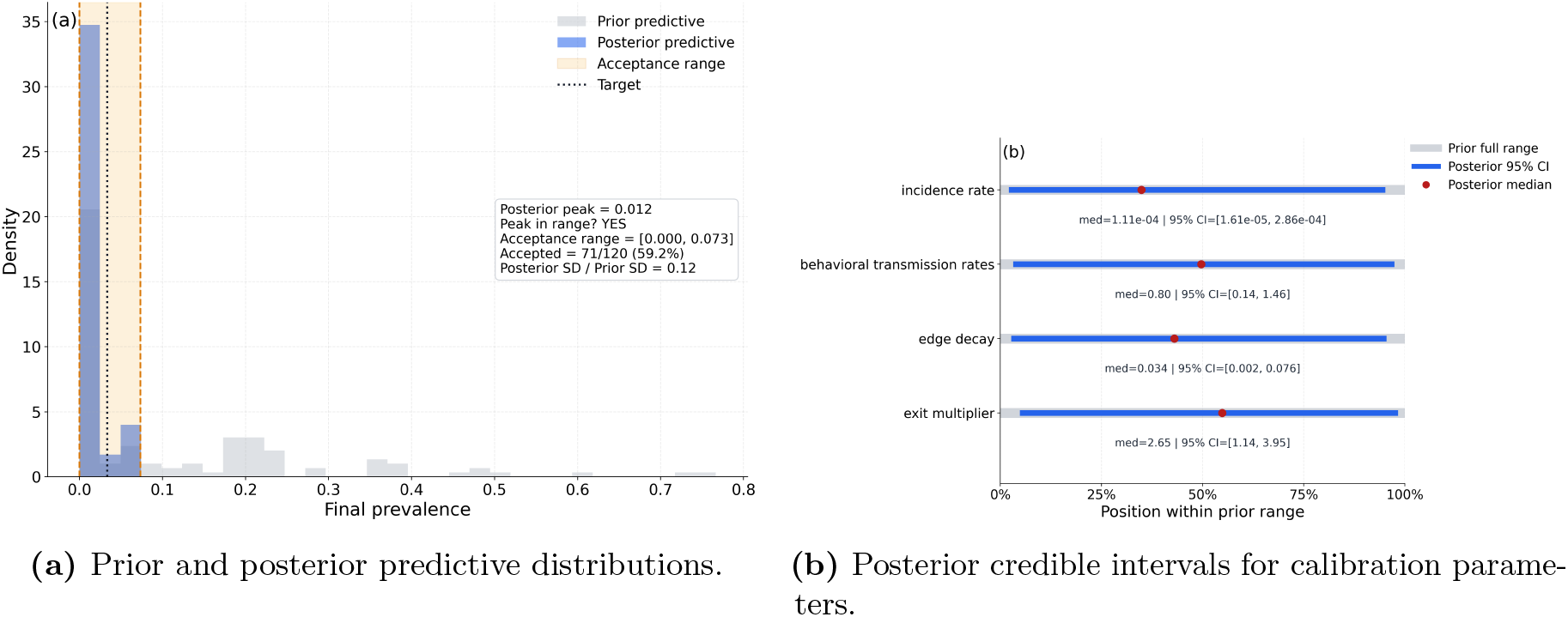
Bayesian calibration under the empirical prevalence target. Panel A shows the prior (gray) and posterior (blue) predictive distributions of the final cardiovascular disease prevalence. The posterior was concentrated within the empirical acceptance region 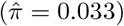, with posterior standard deviation equal to 12% of the prior standard deviation. Panel B presents posterior medians and 95% credible intervals for four calibrated parameters, positioned within their respective prior ranges. Incidence rate showed the strongest concentration toward the lower portion of the prior, consistent with the low target prevalence. Behavioral transmission rates spanned nearly the full prior range, indicating weaker identification under this calibration target.

Parameter-level posteriors showed varying degrees of refinement. The incidence rate posterior median was 1.11 × 10^*−*4^ (95% credible interval 1.6 × 10^*−*5^ to 2.86 × 10^*−*4^), positioned in the lower portion of the prior range and consistent with the low target prevalence. Behavioral transmission rates showed weaker identification, with a posterior median of 0.80 (95% credible interval 0.14–1.46) spanning nearly the full prior range. Edge decay (posterior median 0.034, 95% credible interval 0.002–0.076) and exit multiplier (posterior median 2.65, 95% credible interval 1.14–3.95) showed modest posterior concentrations relative to their priors. These patterns suggest limited but non-zero information gain from a single aggregate prevalence target. They also support the sensitivity analysis reported above as the primary tool for understanding parameter influence on model output.

## 4 Discussion

Coupled dynamics of social connectivity, behavioral adaptation, and demographic renewal jointly shaped chronic disease trajectories in ways that individual risk factor accumulation alone could not produce. Simulations consistently generated lower long-run CVD incidence among highly connected agents. This degree-protective effect arose through two pathways represented in the model: a structural buffering term (*f*_*k*_) through which higher connectivity directly lowered intrinsic risk, and the local behavioral environment that network position created and through which risk propagated dynamically. The present analyses did not separate the relative contributions of these two pathways, and this remains a question for future empirical work. Population turnover acted as a structural dampening mechanism on prevalence, continuously renewing the population with younger healthy agents in a way that no static model could represent. These patterns were robust across a wide parameter space and survived adjustment for all simulated individual-level risk factors.

The degree-protective association in our simulations was consistent with prospective cohort evidence linking social integration to lower cardiovascular risk. Meta-analytic estimates show that social isolation increases coronary heart disease risk by 29% and stroke risk by 32% [5], lack of social support predicts elevated cardiovascular mortality [7], and larger social networks are associated with lower cardiovascular risk over extended follow-up [6, 9–11, 13]. The adjusted HR of 0.43 for the top-tertile degree indicator in our simulations fell within the range of observational estimates that account for confounding, although direct numerical comparison between simulation outputs and empirical effect sizes should be made cautiously given the absence of external validation. Because this degree association persisted after adjustment for the simulated individual-level risk factors, it was consistent with a network-mediated pathway rather than with confounding by the measured covariates alone. We interpreted this as consistent with, rather than proof of, peer influence, since selection and influence remained entangled in the dose-response analysis.

This framework extended prior work on epidemic-network coevolution [28] in three ways. First, CVD was modeled as an absorbing state with no recovery, so that the network permanently loses high-degree nodes to disease and restructures around survivors, unlike SIS and SIR models where recovery allows network restoration. Second, demographic turnover introduces continuous population renewal that produces prevalence equilibria closed-population models cannot generate. Third, behavioral feedback operates through ego-network risk perception rather than global signals, producing local heterogeneity in activity suppression that shapes tie formation and dissolution. Together these features make the framework structurally appropriate for chronic diseases with long latency and age-dependent risk, which acute epidemic models were not designed to represent [34, 35].

This framework also built upon the established tradition of population-level CVD modeling rather than competing with it. Policy models such as the Coronary Heart Disease Policy Model [43] and the IMPACT family of models have shown how much of the decline in CVD mortality reflects risk-factor change versus treatment and have projected the effects of prevention policies [44, 45]. These models treat individuals or subgroups as independent units whose risk factors follow preset paths, with the surrounding social structure held fixed. The present model started from the opposite point: rather than forecasting incidence or testing policy options, it asked how social ties, behavior, and population turnover together produce the risk-factor distributions that those models take as given. It therefore sits at the more mechanistic end of a modeling spectrum that runs from compartmental and life-table models, through microsimulation, to agent-based approaches [25, 26, 32, 33], and it extends that tradition to how social networks and disease evolve together.

The dose-response relationship was consistent with empirical observations of behavioral clustering in social networks. Large cohort studies have documented network clustering of obesity, smoking cessation, and alcohol use [14, 15], with effect sizes suggesting peer influence beyond what shared environment or genetics alone can explain, and agent-based models have shown that behavioral contagion can generate population-level clustering that individual-level mechanisms cannot [46]. Here, “social contagion” refers to behavioral transmission of risk through network ties, not biological infection. Separating survival-model degree effects from the rapid exposure saturation seen in the homophily-contagion regime is something observational studies cannot easily achieve [17], and it represents a primary contribution of the computational approach.

Sensitivity analysis revealed that a small set of parameters accounted for the vast majority of output variability. Baseline incidence rate, activity levels, and behavioral transmission rates jointly explained 78% of variance, while edge decay, similarity weight, and memory decay contributed negligibly. This parsimonious structure identifies the parameters that future empirical work most needs to constrain. Estimates of social activity suppression in response to perceived disease risk and of CVD-based homophily in real social networks would substantially reduce model uncertainty. Bayesian calibration suggested that the model can be anchored to an empirical prevalence target, although behavioral transmission rates remained weakly identified under a single prevalence scalar. Richer calibration data, such as network snapshots paired with longitudinal disease records, would improve parameter identifiability considerably [47].

Several limitations should inform interpretation. As noted, the model is hypothesis-generating rather than predictive, and its quantitative outputs should be interpreted accordingly. Behavioral mechanisms were represented parsimoniously, with risk transmission operating through tie-weighted contact rather than detailed social influence processes such as norm formation or deliberate imitation. The model did not incorporate socioeconomic gradients in network formation, healthcare access, or spatial structure, all of which shape real network-health associations [4, 18]. Generalizability of quantitative estimates requires caution, although the qualitative mechanisms are expected to operate broadly [25, 26, 32, 33].

The model generated four testable predictions for future empirical work. First, longitudinal studies with repeated network measurements should observe lower CVD incidence among individuals with larger ego networks after adjustment for socioeconomic status and baseline health [9, 13]. Second, interventions increasing social connectivity among isolated individuals may alter long-term cardiovascular risk trajectories [48, 49]. Third, homophily effects should be stronger during earlier stages of disease accumulation than later, once saturation equalizes local exposure across connectivity levels. Fourth, the degree-protective association should be most pronounced among individuals with moderate baseline risk, where network-mediated buffering has room to operate. Testing these predictions requires longitudinal datasets pairing repeated network measurements with chronic disease incidence and behavioral risk factors [18, 20]; future extensions could add healthcare network structure [24], cascade effects through alters [19], and temporal network methods [50–52].

Chronic diseases have long been understood through the lens of individual biology and behavior. This framework suggests a complementary perspective: the social structures in which people are embedded and the demographic processes that renew populations are active determinants of disease trajectory rather than background context [20, 53–60]. As longitudinal network data and computational methods become more accessible, frameworks of this kind may help bridge the gap between social epidemiology and the mechanistic traditions of mathematical biology and network science.

## Ethics

This work did not involve human participants, animal subjects, or identifiable personal data. Ethical approval was not required.

## Data accessibility

Python code implementing the agent-based model and all analysis scripts have been deposited in a public repository. The persistent identifier is withheld during the double-anonymous review to ensure author blinding and is provided on the separate title page. Model parameters are documented in Supplementary Table S1.

## 5 Supplementary Material

### 5.1 Supplementary Figure S1: Network degree evolution

The network degree evolution is illustrated in Figure S1. The mean degree declined gradually from an initial value of 9.1, reaching approximately 3.4 over 150 months. This trajectory reflects a regime in which stochastic edge dissolution outpaced compensatory formation driven by homophily and triadic closure. The gradual decline indicates that the network remained structurally coherent throughout the primary simulation analyses.

**Figure S1:**
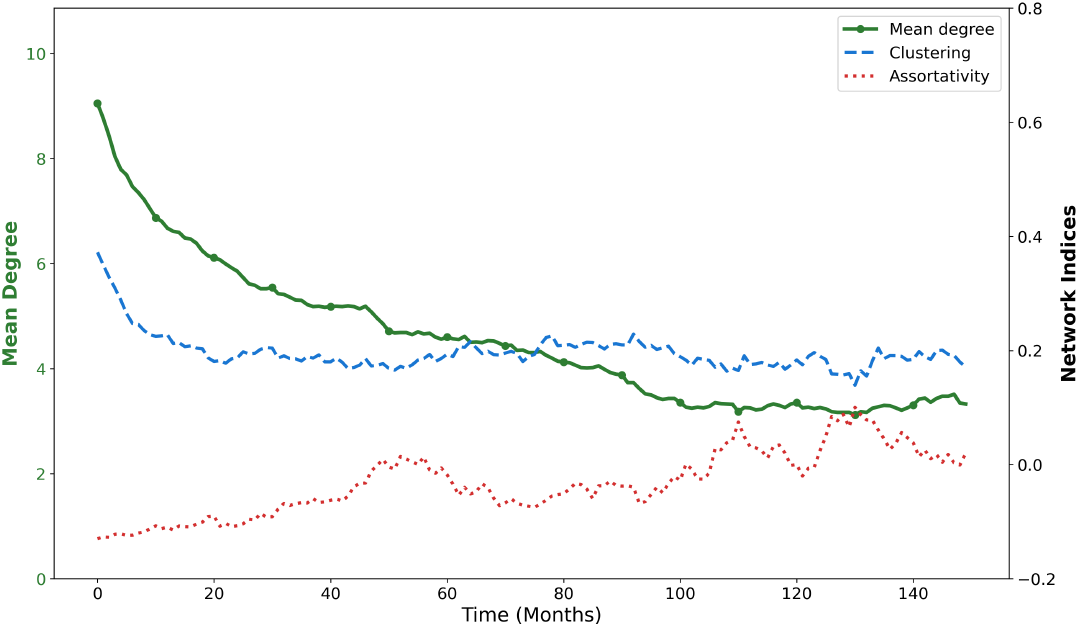
Network degree evolution. Mean network degree declined from an initial value of 9.1, reaching approximately 3.4 over 150 months. The clustering coefficient and assortativity are shown on the secondary axis. This trajectory reflects the balance between stochastic edge dissolution and compensatory formation driven by homophily and triadic closure.

### 5.2 Supplementary Figure S2: Behavioral feedback dynamics

Figure S2 documents two features of the behavioral feedback mechanism. Panel A shows the relationship between population CVD prevalence and mean social activity over time, using simulations in which ego-network weighting parameter *ω*_*i*_ was fixed at 1.0 so that perceived prevalence depended entirely on the local neighborhood. As simulated CVD prevalence rose sharply to a peak of approximately 62% over the first 15 months and then stabilized near 53%, mean activity on the right axis declined from an initial level near 0.14 to a nadir of approximately 0.066, subsequently stabilizing near 0.10, producing a pooled Pearson correlation of *r* = *−*0.771 between prevalence and activity. These patterns are consistent with the behavioral suppression mechanism specified in the model: rising local disease exposure reduces social participation, which in turn reduces transmission opportunities.

Panel B documents the dose-response structure of the social transmission component. Individual CVD probability increased monotonically with both the number of CVD-positive neighbors and weighted infected contact frequency. The relationship with neighbor count followed a saturating logarithmic form, consistent with the functional form specified in the main text, while the relationship with contact frequency showed approximately linear scaling. Spearman rank correlations on the raw event-level data were 0.71 for both exposure metrics (*p <* 0.001). These patterns confirm that exposure breadth and exposure intensity both contribute independently to simulated transmission risk, justifying the two-channel structure of the social-transmission component.

**Figure S2:**
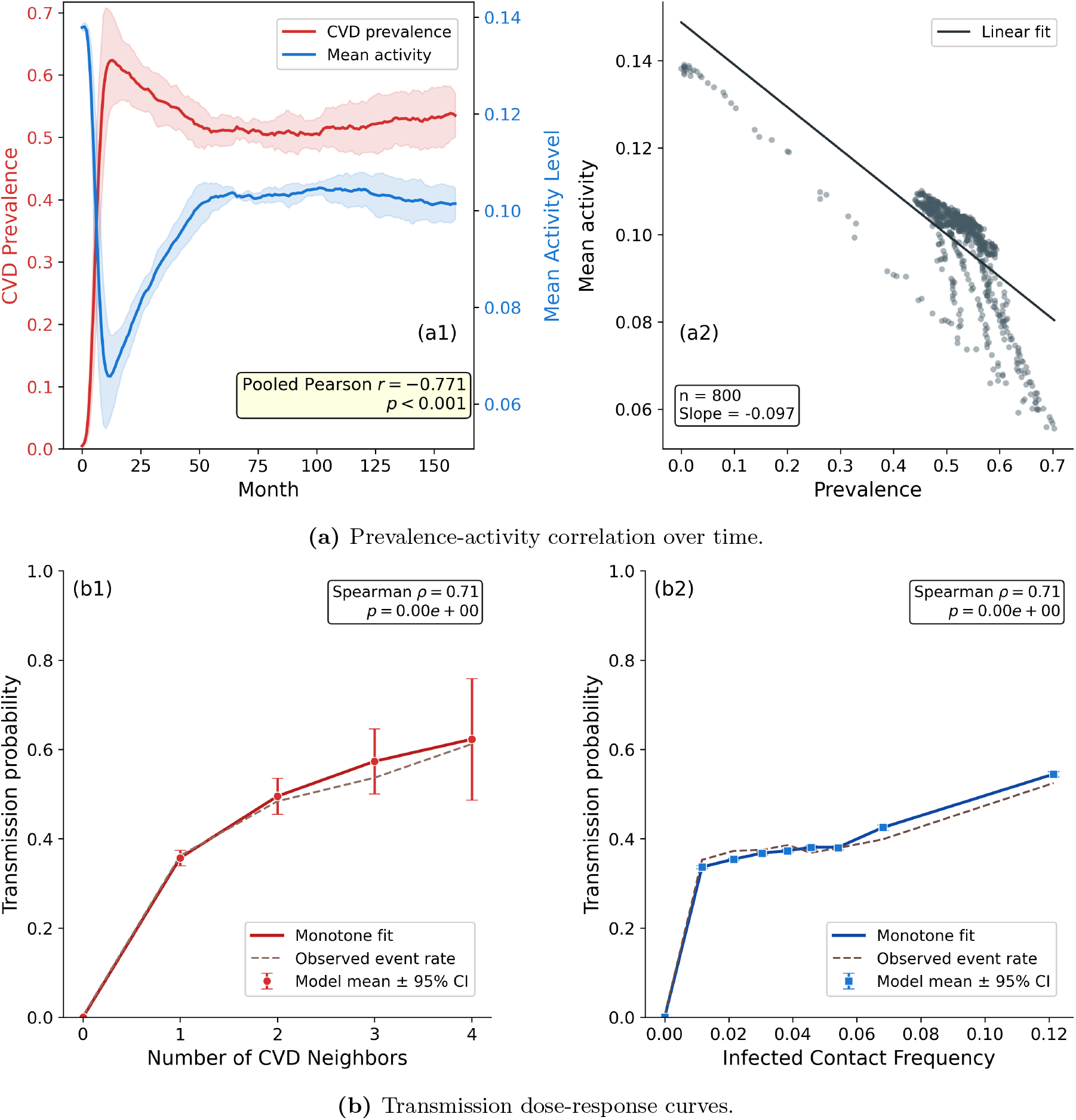
Behavioral feedback and transmission dose-response relationships. Panel A shows the negative correlation between cardiovascular disease (CVD) prevalence and mean activity level (pooled Pearson *r* = *−*0.771, *p* < 0.001) when ego-network weighting was fixed at 1.0. As CVD prevalence rose to a peak of approximately 62% before stabilizing near 53%, mean activity (right axis) declined from approximately 0.14 to a nadir near 0.066, subsequently stabilizing near 0.10, consistent with the behavioral suppression mechanism specified in the main text. The scatter plot (right) was consistent with a negative linear relationship across monthly snapshots (slope = *−*0.097, *n* = 800). Panel B shows individual CVD probability as a function of the number of CVD-positive neighbors (left, saturating logarithmic form, Spearman *ρ* = 0.71, and *p* < 0.001) and weighted infected contact frequency (right, approximately linear scaling, Spearman *ρ* = 0.71, and *p <* 0.001). Both exposure breadth and exposure intensity contributed independently to simulated transmission risk.

### 5.3 Supplementary Figure S3: Parameter space exploration

Figure S3 presents two-dimensional sweeps across parameter pairs that govern network connectivity and behavioral suppression. Panel A shows the final CVD prevalence as a joint function of edge persistence probability *π*_edge_ and network formation scale *σ*. Prevalence increased with both parameters across the grid, reflecting the fact that denser, more persistent networks sustain greater social transmission. The effect of formation scale was stronger at high edge persistence, indicating a degree of interaction between the two network parameters. The dashed green box marks the realistic parameter region used for the primary analyses.

Panel B shows final CVD prevalence as a joint function of behavioral feedback strength *ϕ* and CVD exit multiplier *ξ*. Increasing either parameter reduced simulated prevalence but through distinct mechanisms: higher *ϕ* suppressed social activity and thereby reduced transmission opportunities, while higher *ξ* accelerated the replacement of CVD-positive agents by healthy young entrants. The effects were partially interactive, with strong behavioral suppression reducing the marginal impact of exit multiplier at high *ϕ* values. Together, the two panels confirmed that the model produces internally consistent prevalence surfaces across a wide parameter space, and the primary simulation settings fall within a region of plausible and stable model behavior.

**Figure S3:**
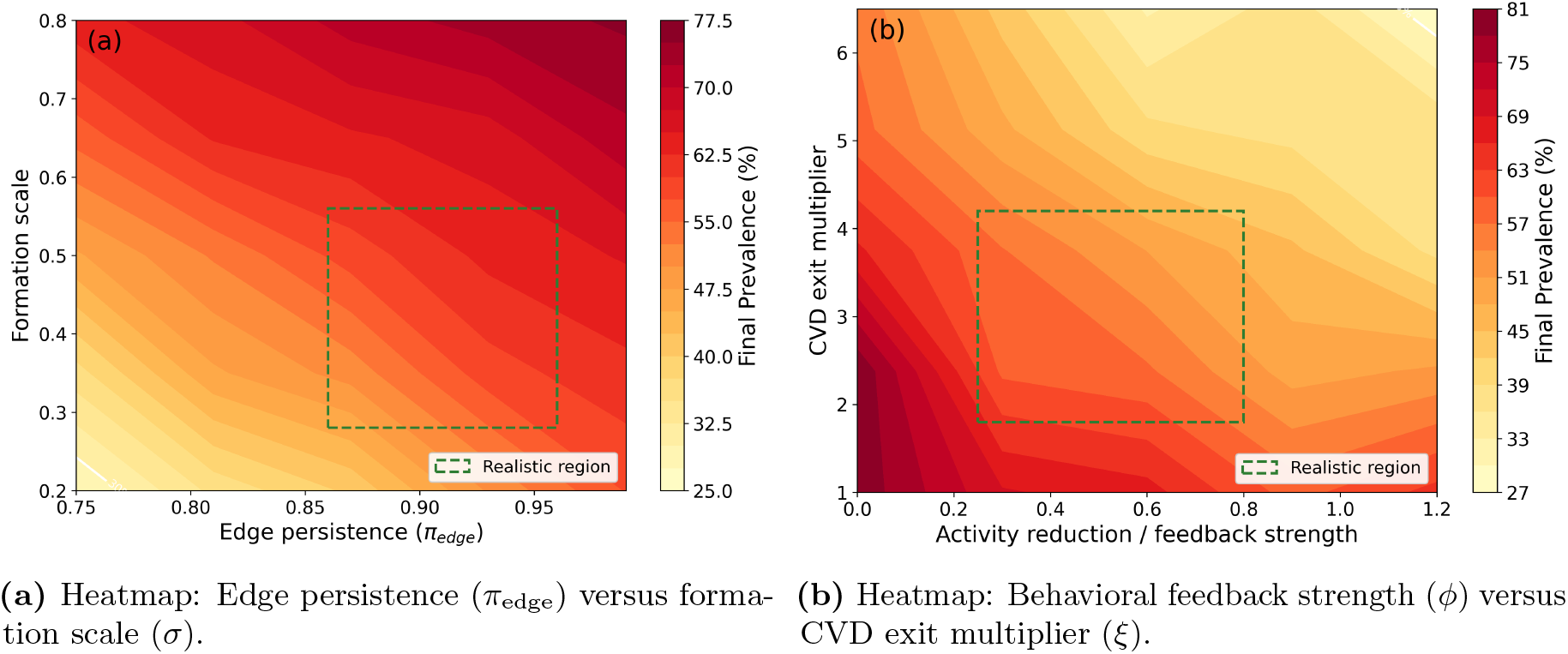
Two-dimensional parameter space exploration. Panel A shows final cardiovascular disease (CVD) prevalence across an 11 × 11 grid of edge persistence probability (*π*_edge_ *∈* [0.75, 0.99]) and network formation scale (*σ ∈* [0.20, 0.80]). Higher values of both parameters increased long-run prevalence by sustaining connectivity and transmission opportunities. Panel B shows the final CVD prevalence across an 11 × 11 grid of behavioral feedback strength (*ϕ ∈* [0.0, 1.2]) and CVD exit multiplier (*ξ ∈* [1.0, 6.5]). Increasing either parameter reduced prevalence through distinct mechanisms: *ϕ* suppressed social activity and reduced transmission, while *ξ* accelerated renewal of the population with healthy young entrants. The effects were nonlinear and partially interactive. Dashed green boxes indicate the realistic parameter regions corresponding to the primary simulation settings.

### 5.4 Supplementary Table S1: Figure-specific parameter settings

Table 1 documents the parameter configuration used for each figure in the main text and each supplementary figure. Because the analyses addressed distinct questions at different scales, parameter settings varied across figures to isolate specific mechanisms or to operate in signal-rich regimes appropriate for the sensitivity analysis and calibration. Population size ranged from *N* = 50 in the compact configurations used for global sensitivity analysis and Bayesian calibration to *N* = 420 in the survival analyses. Simulation horizons ranged from 18 months in the calibration analyses to 240 months in the population-dynamics analyses. All other aspects of the model architecture and monthly update sequence were identical across analyses. Parameters listed as default took the values specified in the `NetworkSimulation` constructor described in the main Methods section. Parameters listed as sampled were varied over the stated range using Latin hypercube sampling, as described in the Global Sensitivity Analysis and Bayesian Calibration subsections of the main text.

**Table 1.**
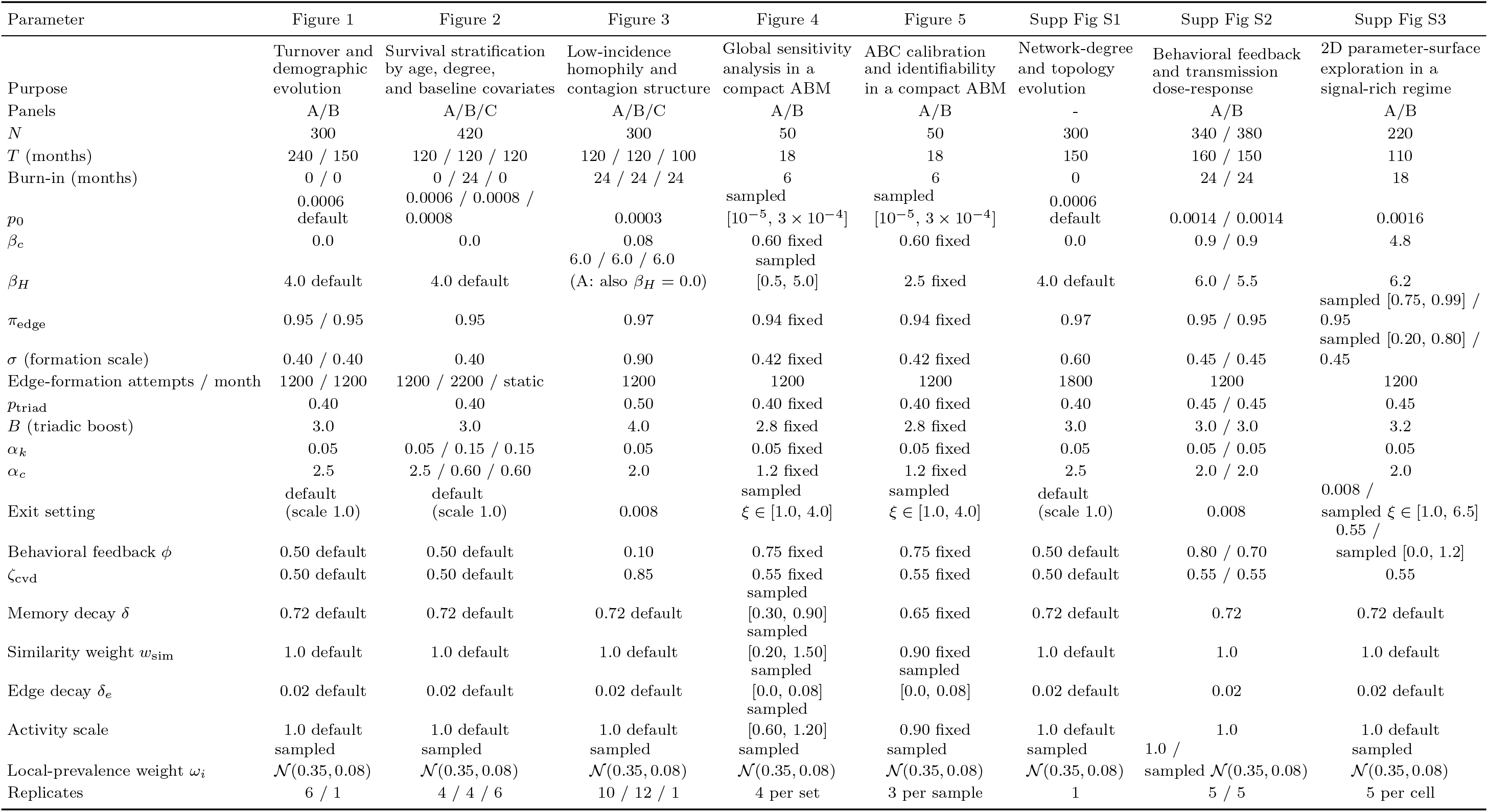
Revised Supplementary Table S1. Figure-specific parameter settings. Figure 1C from the previous table was separated into Supplementary Figure S1. Panel-specific values are listed in panel order (A/B/C or A/B). “Default” denotes the native `NetworkSimulation` constructor defaults. “Sampled” denotes parameters varied over the stated range.

## Notes

### Competing Interest Statement

The authors have declared no competing interest.

